# Therapeutic activity of an inhaled potent SARS-CoV-2 neutralizing human monoclonal antibody in hamsters

**DOI:** 10.1101/2020.10.14.339150

**Authors:** Michael S. Piepenbrink, Jun-Gyu Park, Fatai S. Oladunni, Ashlesha Deshpande, Madhubanti Basu, Sanghita Sarkar, Andreas Loos, Jennifer Woo, Phillip Lovalenti, Derek Sloan, Chengjin Ye, Kevin Chiem, Nathaniel B. Erdmann, Paul A. Goepfert, Vu L. Truong, Mark R. Walter, Luis Martinez-Sobrido, James J. Kobie

## Abstract

SARS-CoV-2 infection results in viral burden in the upper and lower respiratory tract, enabling transmission and often leading to substantial lung pathology. Delivering the antiviral treatment directly to the lungs has the potential to improve lung bioavailability and dosing efficiency. As the SARS-CoV-2 Receptor Binding Domain (RBD) of the Spike (S) is increasingly deemed to be a clinically validated target, RBD-specific B cells were isolated from patients following SARS-CoV-2 infection to derive a panel of fully human monoclonal antibodies (hmAbs) that potently neutralize SARS-CoV-2. The most potent hmAb, 1212C2 was derived from an IgM memory B cell, has high affinity for SARS-CoV-2 RBD which enables its direct inhibition of RBD binding to ACE2. The 1212C2 hmAb exhibits *in vivo* prophylactic and therapeutic activity against SARS-CoV-2 in hamsters when delivered intraperitoneally, achieving a meaningful reduction in upper and lower respiratory viral burden and lung pathology. Furthermore, liquid nebulized inhale treatment of SARS-CoV-2 infected hamsters with as low as 0.6 mg/kg of inhaled dose, corresponding to approximately 0.03 mg/kg of lung deposited dose, mediated a reduction in respiratory viral burden that is below the detection limit, and mitigated lung pathology. The therapeutic efficacy achieved at an exceedingly low-dose of inhaled 1212C2 supports the rationale for local lung delivery and achieving dose-sparing benefits as compared to the conventional parenteral route of administration. Taken together, these results warrant an accelerated clinical development of 1212C2 hmAb formulated and delivered via inhalation for the prevention and treatment of SARS-CoV-2 infection.

## Introduction

The SARS-CoV-2 global pandemic has infected over thirty million people and has resulted in over one million deaths so far. It is expected that new infections will continue for many more months, and the virus will persist endemically for years. Although severe infections and deaths have been reported in all ages and demographics, those over 65 years old and those with pre-existing conditions are at highest risk for death (*1*–*5*). Thus far, no drug or vaccine has been approved by the FDA for the treatment or prevention of SARS-CoV-2 infection. Moreover in the US, the number of COVID-19 patients that are hospitalized represent a small fraction (~10%) of the active cases, which implies that the vast majority of COVID-19 symptomatic patients are not hospitalized and need treatment (*6*). Therefore, facilitating greater treatment coverage will be of importance to control transmissibility and healthcare burden.

Neutralizing antibodies (NAbs) induced either by natural infection or vaccination are likely to be critical for protection from SARS-CoV-2 infection and have been correlated with protection from SARS-CoV-2 in animal studies (*7*–*9*), and passive transfer of neutralizing mAbs has demonstrated prophylactic and therapeutic activity against SARS-CoV-2 infection (*10*–*12*). Emerging results in humans treated with convalescent plasma with high titer NAbs suggest therapeutic activity (*13*, *14*). The primary target for SARS-CoV-2 neutralizing antibodies is the Receptor Binding Domain (RBD), whereby antibodies are expected to inhibit the binding of the SARS-CoV-2 Spike (S) protein to the host Angiotensin Converting Enzyme 2 (ACE2), preventing viral attachment. Already, several human monoclonal antibodies (hmAbs) have been isolated from patients following SARS-CoV-2 infection that are specific for RBD, neutralize SARS-CoV-2, and have anti-viral activity in animal models (*10*, *15*). RBD is used predominantly as the target in clinical stage vaccines and antibody candidates, with preliminary positive clinical responses reported (*6*, *16*).

To obtain more precise resolution of the RBD-specific NAb response, a panel of RBD specific hmAbs were isolated and their molecular features, reactivity profiles and *in vitro* and *in vivo* anti-viral activities were defined. Several high affinity hmAbs with modest somatic hypermutation and potent SARS-CoV-2 neutralizing activity were isolated from IgG, IgA, and IgM memory B cells. In this report, we show that 1212C2 hmAb demonstrated substantial prophylactic and therapeutic activity in hamsters when delivered parenterally. Moreover, when delivered as inhaled liquid aerosols using a commercially available nebulizer, 1212C2 mediated eradication of lung viral load at a substantially higher dosing efficiency than the parenteral route of administration. The combination of a potent SARS-CoV-2 mAb and inhaled, local lung delivery using widely available nebulizers could provide for a promising treatment option.

## Results

### Isolation of RBD+ B cells and hmAb screening

To identify and isolate RBD-specific B cells, recombinant RBD protein was expressed, biotinylated, and used to form streptavidin-conjugated RBD tetramers to various fluorochromes. Peripheral blood CD27+ memory B cells binding RBD were single cell sorted from convalescent SARS-CoV-2 patients and the immunoglobulin heavy and light chain variable regions of the B cells cloned to generate IgG1 recombinant hmAbs (**Fig 1A**). Twenty hmAbs were isolated that exhibited substantial binding to SARS-CoV-2 RBD. These hmAbs all bound to recombinant SARS-CoV-2 RBD and S1 D614G proteins and exhibited varying reactivity SARS-CoV-2 S1S2 protein (**Fig 1B**). As expected, minimal to no reactivity was observed to SARS-CoV-2 Nucleocapsid (N) protein or HepG2 cell lysate for most hmAbs, indicating high specificity for SARS-CoV-2 RBD and minimal off-target binding. REGN10987, REGN10933, and CB6/JS016 hmAbs were synthesized and included as positive controls (*15*, *17*). To ascertain the avidity of the hmAbs for SARS-CoV-2 RBD, their binding stability was determined in the presence of the chaotropic agent 8M urea. The hmAbs 1206D1, 1212C2, 1212F5, and 1215D1 retained at least 50% of their binding activity to SARS-CoV-2 RBD and S protein (**Fig 1C**). Binding affinity to RBD was further tested for a subset of the hmAbs by surface plasmon resonance (SPR), demonstrating a variety of high affinity hmAbs exhibiting equilibrium dissociation constants (*K*D) ranging from 77 pM to ~9 nM (**Fig 1D**). Several of the hmAbs (1212C2, 1215D1, 1215D5) exhibited ~10-fold higher affinity to RBD than control hmAbs (REGN10987, REGN10933 (*17*), and CB6/JS016 (*15*)), predominantly due to their slower off rates (*k*d). These results indicate that this panel of hmAbs recognize SARS-CoV-2 RBD specifically and with high affinity.

**Figure 1.**
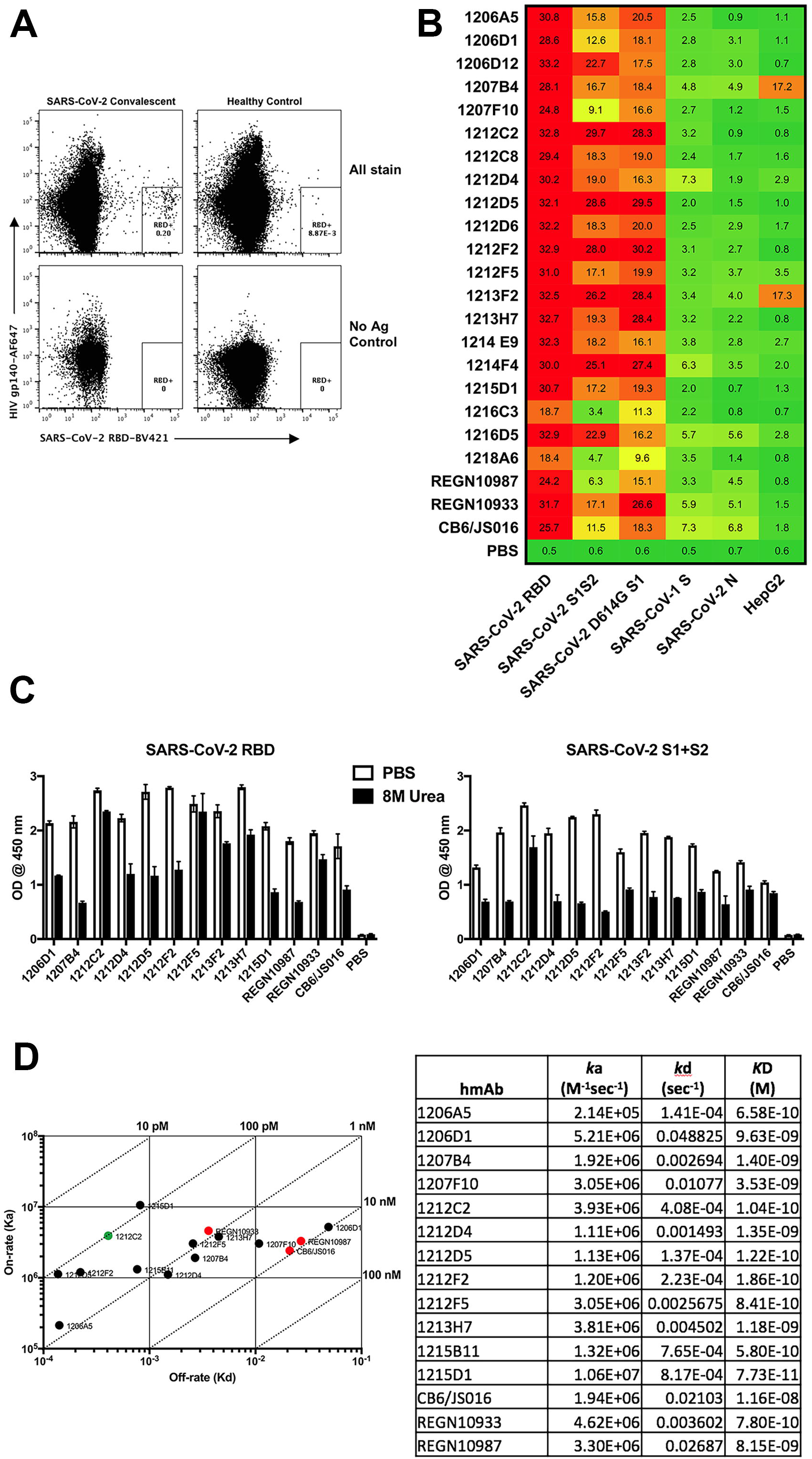
Isolation of SARS-CoV-2 RBD specific human monoclonal antibodies. (**A**) Representative gating strategy for RBD+ B cell isolation. Initial plots are gated on live CD3- CD4- CD19+ Annexin V-CD27+ B cells. (**B**) hmAbs were tested at 10 and 1 μg/ml by ELISA for binding to indicated protein, Area Under the Curve (AUC) is indicated. (**C**) hmAbs were tested at 10 μg/ml for binding to indicated protein by ELISA in the presence or absence of 8M urea treatment. (D) Binding to SARS-CoV-2 RBD was determined by surface plasmon resonance.

### Recognition and neutralization of SARS-CoV-2

The process of entry into a susceptible host cell is an important determinant of infectivity and pathogenesis of viruses, including coronaviruses (*18*, *19*). SARS-CoV-2 relies on the ability of its S glycoprotein to bind to the ACE2 receptor through its RBD driving a conformational change that culminates in the fusion of the viral envelope with the host cell membrane, and cell entry (*20*). The hmAbs were tested for neutralization of live SARS-CoV-2 using a virus plaque reduction microneutralization (PRMNT) assay we previously described (*21*), with the hmAb and virus pre-incubated prior to culture with susceptible Vero E6 cells (pre-treatment) or allowing virus adsorption to Vero E6 cells to occur for 1 hour prior to addition of the hmAb (post-treatment). This gives an opportunity for the virus to initiate viral entry by binding to the cell surface receptor, potentially distinguishing the hmAb’s ability to preferentially block later steps of virus entry into the cell or by inhibiting the cell-to-cell spread of virus progeny. The panel of hmAbs neutralized SARS-CoV-2 in both pre- and post-treatment conditions at NT_50_ of 1 μg/ml or less, with 1212F5, 1212C2, and 1213H7 exhibiting the highest potency, with NT_50_ of 100 ng/ml or less (**Fig 2A**). The hmAb 1215D1 was more effective in neutralizing in post-treatment (NT_50_ = 59 ng/ml) compared to pre-treatment (NT50 = 226 ng/ml)). The 1212C2 hmAb was particularly potent (pre NT_50_ = 45 ng/ml, post NT_50_ = 32 ng/ml) (**Fig 2B**). Testing of 1212C2 in a SARS-CoV-2 VSV vectored pseudovirus assay confirmed its potent neutralizing activity (NT_50_ = 1.9 ng/ml) that was similar to REGN10987 and CB6/JS016 (**Supplemental Table 1**).

**Figure 2.**
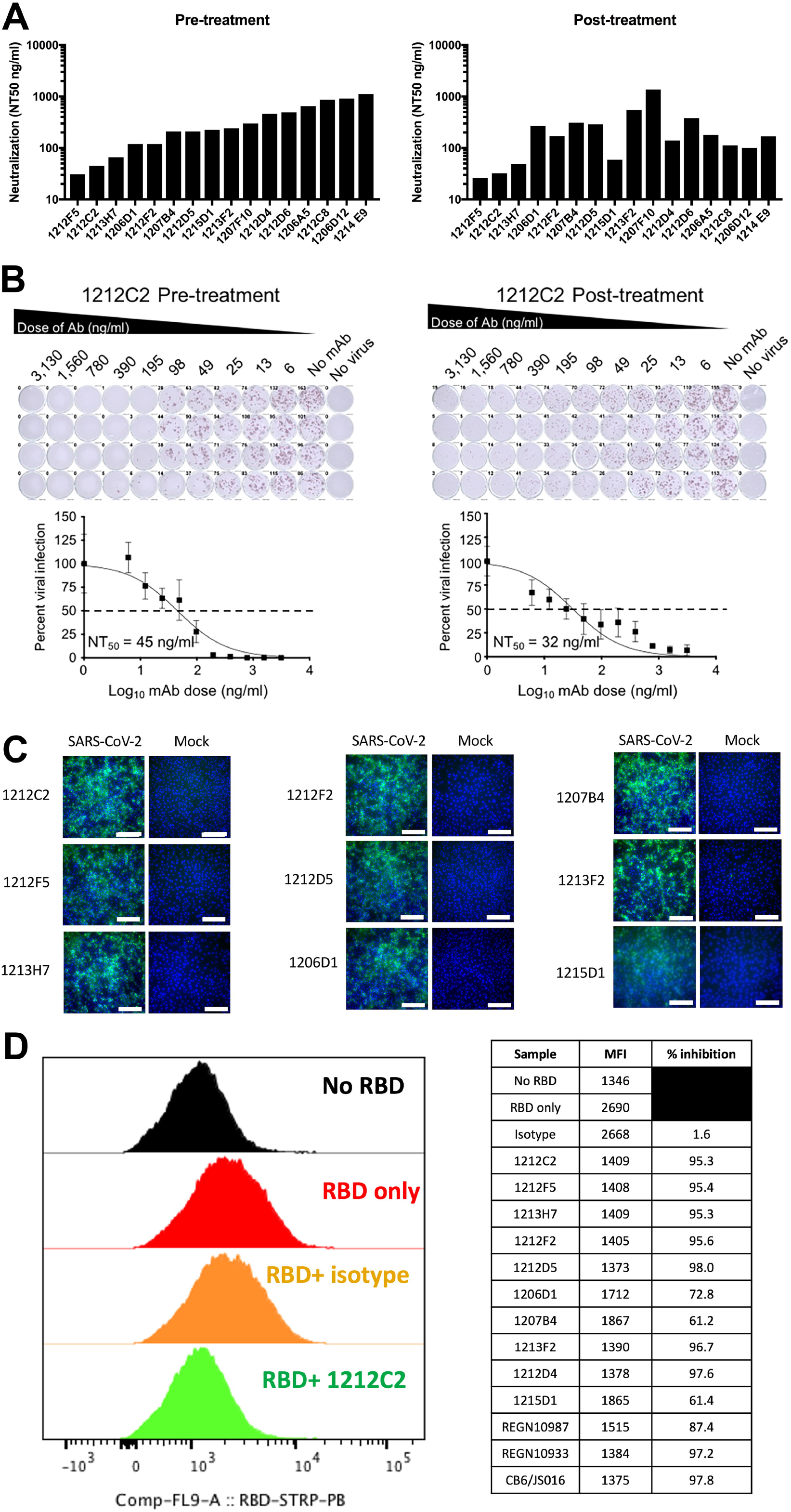
Neutralization of SARS-CoV-2 infection and RBD binding to ACE2. (**A**) Indicated hmAbs were incubated with live SARS-CoV-2 (100 PFU/well) 1 hour prior to (pre-treatment) or 1 hour after (post-treatment) addition to Vero E6 cells. hmAbs were tested in quadruplicate cultures and NT_50_ indicated. (**B**) Representative titration curve of 1212C2 hmAb presented. (**C**) Binding of indicated hmAb to SARS-CoV-2 or mock infected Vero E6 cells measured by immunofluorescence. (D) Indicated hmAb was incubated with recombinant biotinylated RBD-protein prior to incubation with HEK293-ACE2 cells measured by flow cytometry. Plot gated on 7AAD-ACE2+ cells.

The panel of hmAbs clearly recognized SARS-CoV-2-infected Vero E6 cells as evident by immunofluorescence (**Fig 2C and Supplemental Figure 1**). No background staining of mock-infected cells was evident with any of the hmAbs, consistent with their high affinity specificity for SARS-CoV-2 S. The hmAbs also exhibited binding to SARS-CoV-2 viral lysate (**Supplemental Figure 2**). The ability of the hmAbs to directly inhibit the binding of RBD to ACE2 was determined using HEK293 cells overexpressing ACE2. All of the tested RBD-specific hmAbs inhibited the binding of recombinant SARS-CoV-2 RBD protein to the ACE2 expressing cells (**Fig 2D**). Inhibition was nearly complete with the exceptions of 1206D1, 1207B4, and 1215D1. These results demonstrate the potent *in vitro* neutralizing and binding activity of these SARS-CoV-2 RBD specific hmAbs.

### Molecular features and clonal dynamics

The most potent hmAbs, 1212C2 and 1212F5 were both isolated from IgM+ B cells, belong to the same clonal lineage that utilizes the VH1-2 heavy chain gene, and exhibited modest somatic hypermutation, with 1212C2 further mutated from germline compared to 1212F5 (8.2% vs 6.1% amino acid VH) (**Table 1**). Most of the hmAbs were isolated from IgG1 expressing B cells, while 1215D1 and 1212C8 were isolated from IgA expressing B cells. All hmAbs exhibited somatic hypermutation (2.0%-9.1% VH) suggesting they arose from multiple rounds of germinal center reactions. VH3-66 and Vk1-9 gene usage was dominant among the hmAbs. Targeted VH-deep sequencing of the 1212C2/1212F5 clonal lineage from contemporary peripheral blood B cells identified numerous members (**Fig 3**). The lineage was dominated by IgM and IgG expressing B cells, with several expanded nodes of identical clones being expressed by both IgM and IgG expressing B cells.

**Table 1.** Molecular Characteristics of hmAbs

| hmAb | VH | DH | JH | Isotype | %<br>mutation<br>AA | VL | JL | %<br>mutation<br>AA |
| --- | --- | --- | --- | --- | --- | --- | --- | --- |
| **1206D1** | IGHV1-2 | IGHD6-13 | IGHJ3 | IgG1 | 8.2 | IGLV2-8 | IGLJ2 | 8.2 |
| **1207F10** | IGHV1-2 | IGHD5-12 | IGHJ4 | IgG3 | 3.1 | IGLV6-57 | IGLJ3 | 4.1 |
| **1212C2** | IGHV1-2 | IGHD5-18 | IGHJ4 | IgM | 8.2 | IGLV2-23 | IGLJ2 | 5.2 |
| **1212F5** | IGHV1-2 | IGHD5-18 | IGHJ4 | IgM | 6.1 | IGLV2-23 | IGLJ2 | 3.1 |
| **1213H7** | IGHV1-58 | IGHD2-15 | IGHJ3 | IgG1 | 7.1 | IGKV3-20 | IGKJ1 | 3.2 |
| **1206A5** | IGHV3-9 | IGHD1-1 | IGHJ6 | IgG1 | 2.0 | IGLV2-14 | IGLJ2 | 5.1 |
| **1207B4** | IGHV3-9 | IGHD5-12 | IGHJ4 | IgG1 | 7.1 | IGLV1-44 | IGLJ2 | 3.1 |
| **1215D1** | IGHV3-9 | IGHD4-23 | IGHJ6 | IgA1 | 9.1 | IGLV6-57 | IGLJ2 | 3.1 |
| **1206G12** | IGHV3-15 | IGHD3-22 | IGHJ3 | IgG1 | 5.0 | IGKV1-39 | IGKJ4 | 6.3 |
| **1213F2** | IGHV3-53 | IGHD3-16 | IGHJ3 | IgG1 | 8.2 | IGKV1-9 | IGKJ3 | 4.3 |
| **1206D12** | IGHV3-66 | IGHD2-21 | IGHJ6 | IgG1 | 7.2 | IGKV1-33 | IGKJ2 | 2.1 |
| **1212C8** | IGHV3-66 | IGHD5-24 | IGHJ6 | IgA2 | 7.2 | IGKV1-12 | IGKJ4 | 4.3 |
| **1214E9** | IGHV3-66 | IGHD3-16 | IGHJ6 | IgG1 | 7.2 | IGKV1-9 | IGKJ2 | 0 |
| **1212D4** | IGHV3-66 | IGHD3-3 | IGHJ6 | IgG1 | 6.2 | IGKV1-9 | IGKJ2 | 6.4 |
| **1212F2** | IGHV3-66 | IGHD2-15 | IGHJ6 | IgG1 | 4.1 | IGKV1-9 | IGKJ5 | 3.2 |
| **1212D5** | IGHV3-66 | IGHD3-3 | IGHJ3 | IgG1 | 6.2 | IGKV1-5 | IGKJ2 | 4.2 |
| **1212D6** | IGHV3-66 | IGHD3-10 | IGHJ4 | IgG1 | 8.2 | IGKV1-9 | IGKJ3 | 2.1 |

**Figure 3.**
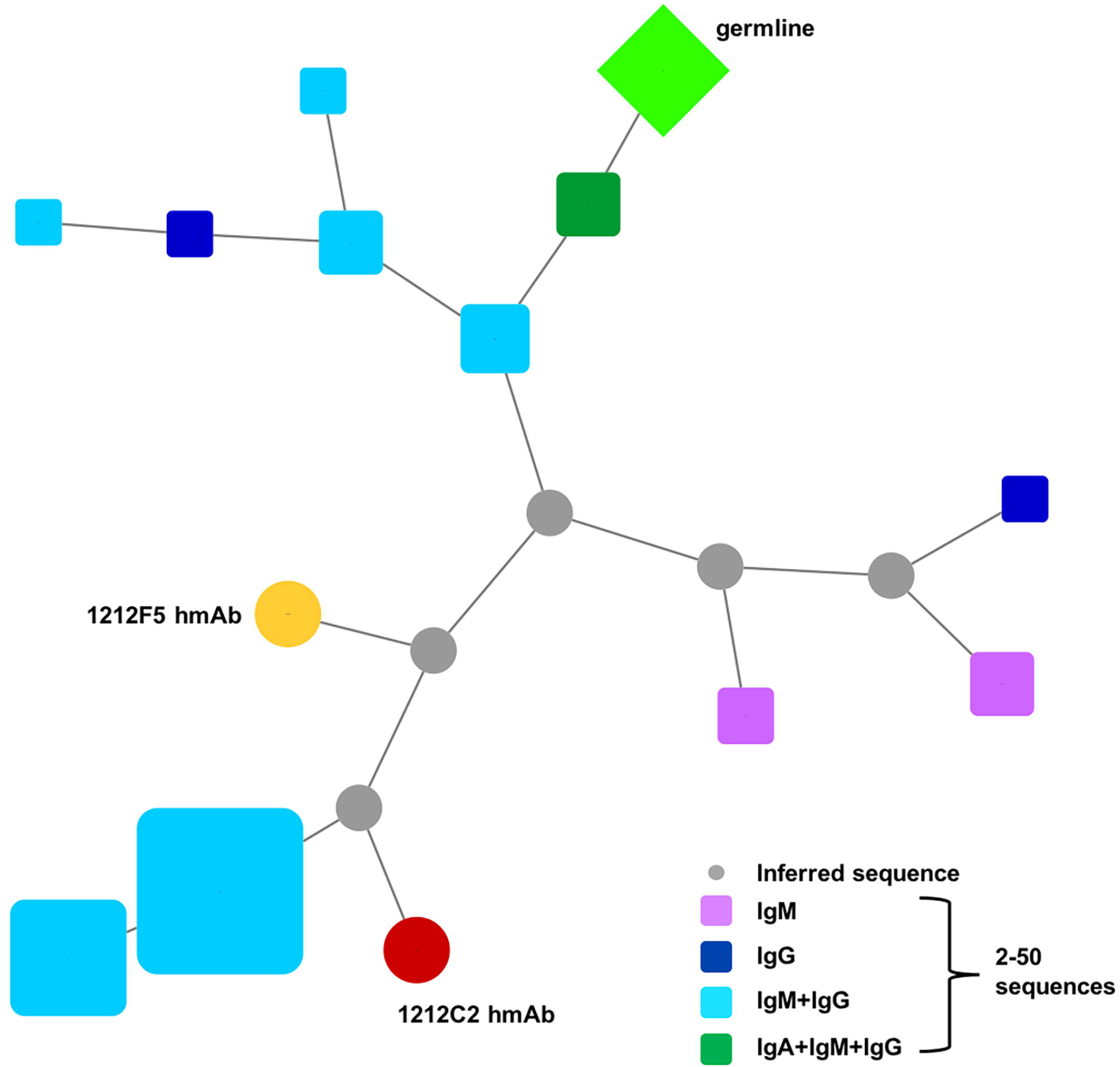
Phylogenic analysis of 1212C2 hmAb lineage. Lineage members defined as same heavy chain V and J gene usage, HCDR3 length, and ≥85% HCDR3 similarity. Germline sequence is represented by green diamond, the 1212C2 hmAb sequence is represented by the red circle, the 1212F5 hmAb sequenced is represented by orange circle. Size of symbols are proportional to the number of identical sequences obtained of an individual lineage member (N□=□2–50), with the exception of the germline and 1212C2 and 1212F5 hmAb sequences. Relationship between lineage members determined by amino acid sequence similarity.

### Epitope Mapping

SPR epitope mapping was performed and used to cluster the hmAbs into five major epitopes (A-E, **Fig 4**). With one exception, all isolated hmAbs are part of the A epitope, that includes 1212C2. Thus, 1212C2 efficiently blocked all hmAbs from binding to RBD, except 1215B11 (E epitope), whose epitope overlaps with CR0322. Within the footprint of the A epitope there are several sub-epitopes. In particular, we highlight hmAbs 1207B4/1215D1 (B-epitope) and 1213H7 (C-epitope) that exhibit essentially no overlap with one another, based on their ability to simultaneously bind RBD at greater than 90% of their expected binding levels. Consistent with their distinct classification, B-epitope hmAbs exhibit a reduced ability to block RBD attachment to ACE2 expressing cells (**Fig 2D**).

**Figure 4.**
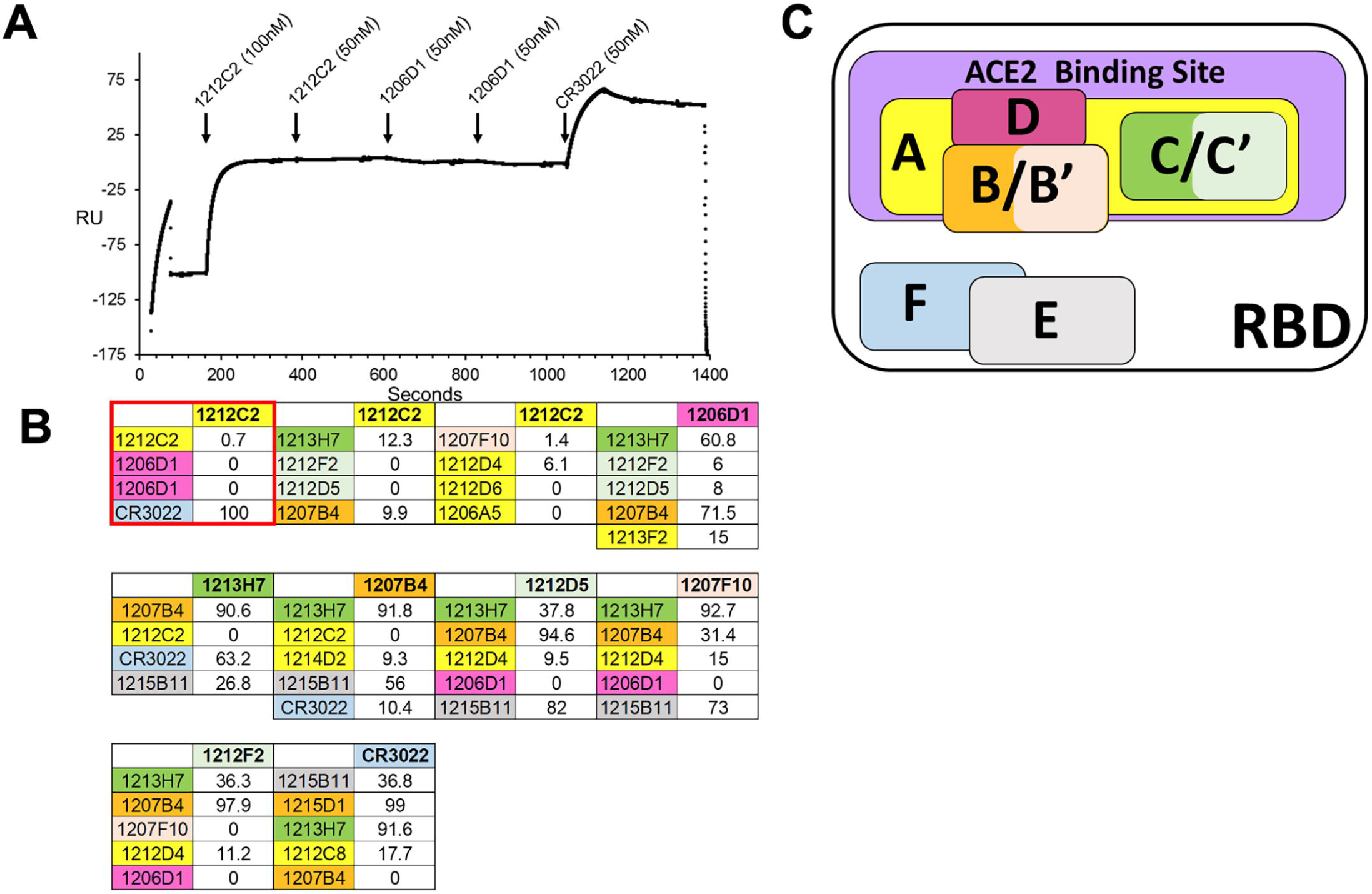
Surface plasmon resonance epitope mapping. (**A**) Representative sensorgram from the SPR competition assays used to subset hmAbs into distinct RBD binding epitopes. For each assay, a series of hmAbs were sequentially injected over immobilized SARS-CoV-2 RBD. In this example, 1212C2 was injected first, followed by a second injection of 1212C2, two injections of 1206D1, and last injection was CR3022. (**B**) Summary of all epitope mapping data, where each block (first experiment from 4A in the red box) with a bold hmAb at the top represents a different experiment (10 experiments total). The bold hmAb is the “first” hmAb injected. The percent binding (100 = 100% binding and 0 = 0% binding) of subsequent hmAbs is recorded. mAbs were considered to have a different epitope (denoted by a distinct color) if they exhibited binding levels >30% in the presence of other mAbs. Thus, in the first experiment, CR3022 is defined as a new epitope (cyan), distinct from 1212C2. (**C**) Schematic diagram of NmAb RBD epitopes defined in the mapping experiment. Five major epitopes (A-E) were identified, where the E epitope overlaps with control mAb CR3022 (cyan, epitope F). Four of the five epitopes (A-D) are located within the ACE2 binding site (purple) and all NmAbs are blocked by the 1212C2 epitope A (yellow). NmAbs with epitopes similar to B (orange) and C (green) are defined as B’ (light orange) and C’ (light green), respectively. The 1212C2 epitope A (yellow) blocks binding of all NmAbs with the exception of 1215D11, that occupies epitope E. Epitopes B and C are also blocked by epitope A NmAbs but exhibit limited competition with each other.

### Prophylactic and therapeutic activity in hamsters

The *in vivo* activity of 1212C2 and 1206D1 was evaluated in the golden Syrian hamster model of SARS-CoV-2 infection (*22*). 1212C2 was chosen based on its potent *in vitro* neutralizing activity and high affinity, while 1206D1 was chosen based on its *in vitro* neutralizing activity and distinct affinity and reactivity profile from 1212C2.

To test for prophylactic activity, 10 mg/kg of hmAb was administered by intraperitoneal (IP) injection 6 hours prior to intranasal (IN) challenge with SARS-CoV-2. At 2 days post infection (dpi), all PBS control and isotype control hamsters had detectable live virus as measured by plaque assay in their nasal turbinate and lungs. In contrast at 2 dpi, hamsters that received 1212C2 already started to exhibit meaningful viral load reduction in their nasal turbinate and lungs (**Fig 5A**). At 4 dpi virus was detected in the nasal turbinates and lungs of all hamsters in the PBS and isotype control groups, although an overall decrease compared to 2 dpi, consistent with the viral dynamics of SARS-CoV-2 infection in hamsters (*22*). In comparison to the control groups at 4 dpi, prophylactically treated 1212C2 hamsters exhibited eradication of viral loads in the nasal turbinates and lungs in 3 of 4 animals. Consistent with the viral load reduction, 1212C2 treated animals exhibited significantly less lung pathology compared to the PBS treated group at 2 dpi and 4dpi, with this reduction also reaching significance compared to the isotype control group at 4 dpi (**Fig 5C**). There was ~80% reduction in lung pathology at 4 dpi when 1212C2 was given prophylactically. 1206D1 hmAb exhibited modest activity, noted by 50% of hamsters having no detectable virus in the nasal turbinate, and all 1206D1 hamsters having detectable virus in the lungs, although trending to lower titers compared to PBS and isotype control groups. These *in vivo* results are consistent with the lower *in vitro* viral neutralization activity of 1206D1 in comparison to 1212C2 (**Fig 2A**). Overall, 1212C2 demonstrated substantial prophylactic activity as evident by sterilizing protection in 63% of hamsters.

**Figure 5.**
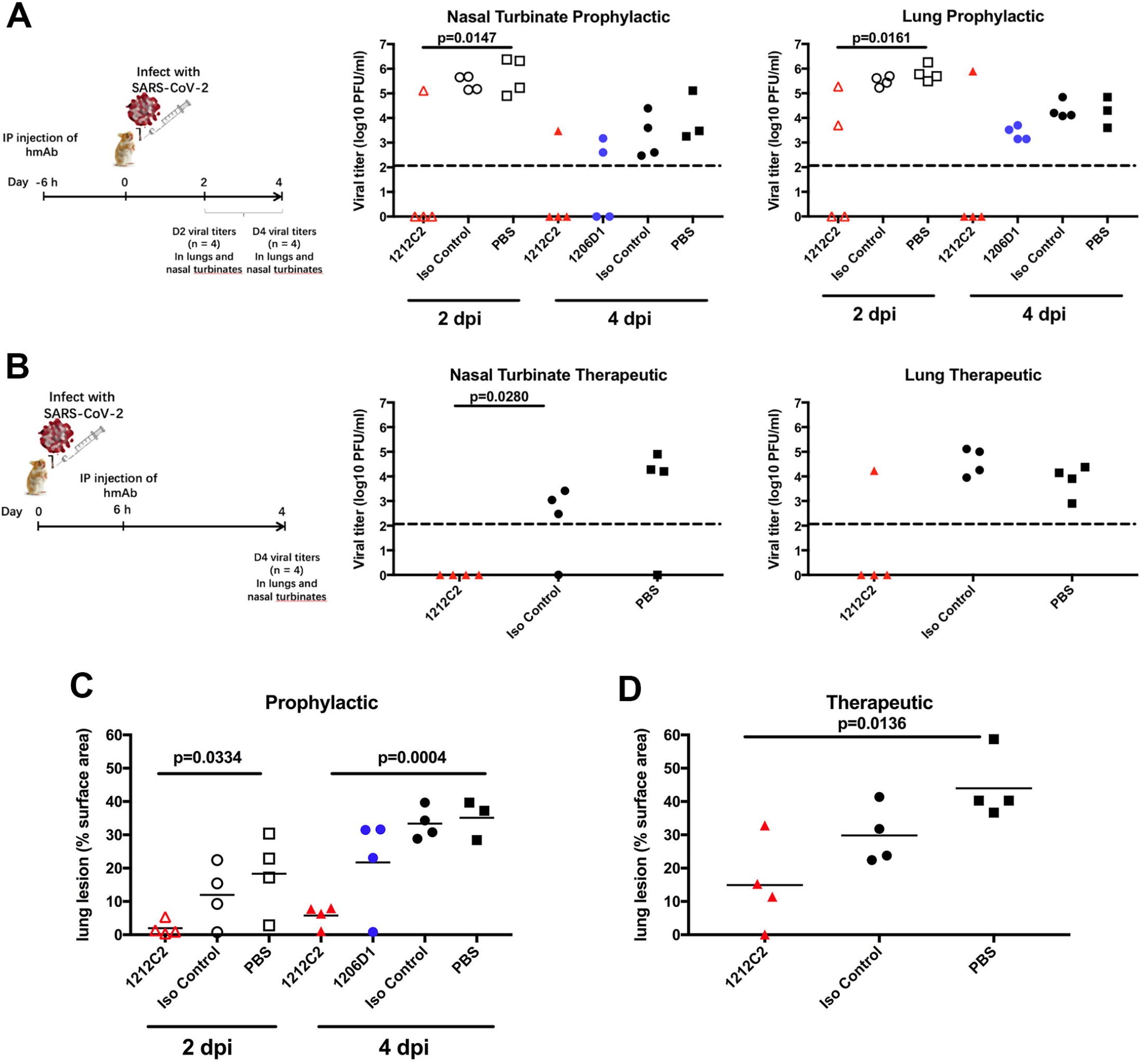
Prophylactic and therapeutic activity of 1212C2 hmAb in SARS-CoV-2 infected hamsters. Golden Syrian hamsters were (**A**) prophylactically treated IP with 10 mg/kg of indicated hmAb or PBS 6 h prior to intranasal challenge with 2×10^5^ PFU SARS-CoV-2 or (**B**) therapeutically treated IP with 25 mg/kg of indicated hmAb or PBS 6 h following intranasal challenge with SARS-CoV-2. Virus present in nasal turbinates and lungs was determined by plaque assay. Dotted line indicates limit of detection. Distribution of pathologic lesion, including consolidation, congestion, and pneumonic lesions were measured using ImageJ and represented as the percent of the total lung surface area in prophylactically treated (**C**) and therapeutically treated (**D**) animals. Each symbol represents an individual animal.

Therapeutic activity was tested by treatment with 25 mg/kg of 1212C2 6 hours following intranasal infection of hamsters with SARS-CoV-2 and evaluating viral burden at 4 dpi only due to limited availability of hmAb. SARS-CoV-2 was detected in the nasal turbinates in 3 of 4 animals in the PBS and isotype control groups. No virus was detected in the nasal turbinates of the 1212C2 treated hamsters. SARS-CoV-2 was detected in the lungs of all PBS and isotype control treated hamsters, but only in 1 of 4 of the 1212C2 treated hamsters (**Fig 5B**). Overall, 1212C2 demonstrated substantial therapeutic activity, reducing virus to undetectable levels in 75% of the treated hamsters.

SARS-CoV-2 infection in hamsters results in gross lung lesions, including consolidation, congestion, and atelectasis (*22*). Therapeutically, significantly less lung pathology was also observed in the 1212C2 treated hamster group compared to the PBS treated group, and ~58% reduction in lung pathology in 1212C2 group compared to the control hamsters (**Fig 5D**). Importantly, these differences observed in lung lesions are consistent with the viral burden seen in the upper and lower respiratory tract (**Fig 5A and 5B**). Overall, these results demonstrate the ability of 1212C2 to substantially reduce viral burden and lung pathology of SARS-CoV-2 infection when used either prophylactically or therapeutically.

### Delivery of inhaled hmAb in hamsters

Therapeutic mAbs are typically delivered systemically through intravenous infusion or intramuscular injection. Such delivery typically results in less than 0.1% of the injected dose in the lung epithelial lining fluid at Cmax, thus highly inefficient in achieving optimal concentrations in the respiratory tract for the protection or treatment of respiratory infections, such as SARS-CoV-2 (*23*).

To evaluate the efficiency of lung delivered dosing and clearance of an IgG1 hmAb when delivered as an IP bolus dose as compared to an inhaled nebulized aerosol, hamsters were administered with the non-specific IgG used in the challenge studies and the serum and bronchoalveolar lavage (BAL) concentrations were analyzed at 30 minutes and 42 hours post-dosing (**Supplemental Table 2**). Intraperitoneal administration of approximately 25 mg/kg of hmAb resulted in high serum concentration but little to no detectable hmAb in the BAL at 30 mins post treatment (**Supplemental Table 2**). However, 42 hours later, the hmAb was detectable in the BAL, suggesting that IP injected hmAb gradually penetrates the lungs from the serum compartment. Inhalation (IH) administration of the hmAb using liquid aerosols delivered whole body to the animals from a commercially available nebulizer (Aerogen Aeroneb Solo nebulizer) showed that approximately 1.7% of the inhaled dose was deposited in the BAL. The low percentage of the inhaled dose that is deposited in the lungs of hamsters is in line with prior lung uptake studies for the approximately 4 μm diameter droplets produced by the nebulizer used here (volume mean diameter measured by laser diffraction 4.1 μm), and is a result of a higher degree of inertial impaction of liquid aerosols in the upper airways of small animals (*24*). Nevertheless, such lung deposited (BAL) dose is still substantially higher at both time points than the BAL concentration achieved with the IP route. A higher lung deposited dose afforded by the inhaled route demonstrates higher delivery efficiency to the lungs than the IP route. At 42 hours post-inhaled dose, the mAb is significantly cleared from the BAL, which appears to be faster clearance rate than an expected lung half-life of ~8 days reported in other studies (45).

### Therapeutic activity of inhaled 1212C2 hmAb in hamsters

The Fc of 1212C2 was modified with the LALA mutation to reduce FcR binding and subsequent Fc-mediated effector functions (*25*), and further modified to increase half-life (*26*) (referred to as ‘1212C2-HLE-LALA’). To determine the efficacy of inhaled 1212C2 and evaluate the therapeutic dose-response, hamsters were infected intranasally with 2×10^5^ PFU SARS-CoV-2 and treated with a single dose of 1212C2 hmAb or isotype control hmAb 12 hours later using IH or IP routes. As measures of efficacy, body weight was recorded, pulmonary lung lesions were measured, and viral titers quantitated from nasal turbinates and lungs on days 2 and 4 following infection.

At 2 dpi virus was detected in the nasal turbinate and lungs of all infected control hamsters (PBS and isotype control hmAb) (**Fig 6A**). Virus was not detectable in the lungs of any hamsters treated via inhalation with 1212C2-HLE-LALA at 16.3 mg/kg and virus was significantly lower in their nasal turbinates compared to controls. In contrast, virus was detectable at 2 dpi in the lungs of 50% of the hamsters treated IP with 1212C2-HLE-LALA at 25 mg/kg, and not significantly decreased in nasal turbinates compared to controls. At 2 dpi, virus was only detectable in the lungs of 25% and 50% of the hamsters treated via inhalation with a lung delivered dose of 3.2 mg/kg and 0.6 mg/kg of 1212C2-HLE-LALA, respectively. At 2 dpi, 16.3 mg/kg and 3.2 mg/kg inhaled 1212C2-HLE-LALA was superior in reducing viral titer in the lungs compared to IP administration.

**Figure 6.**
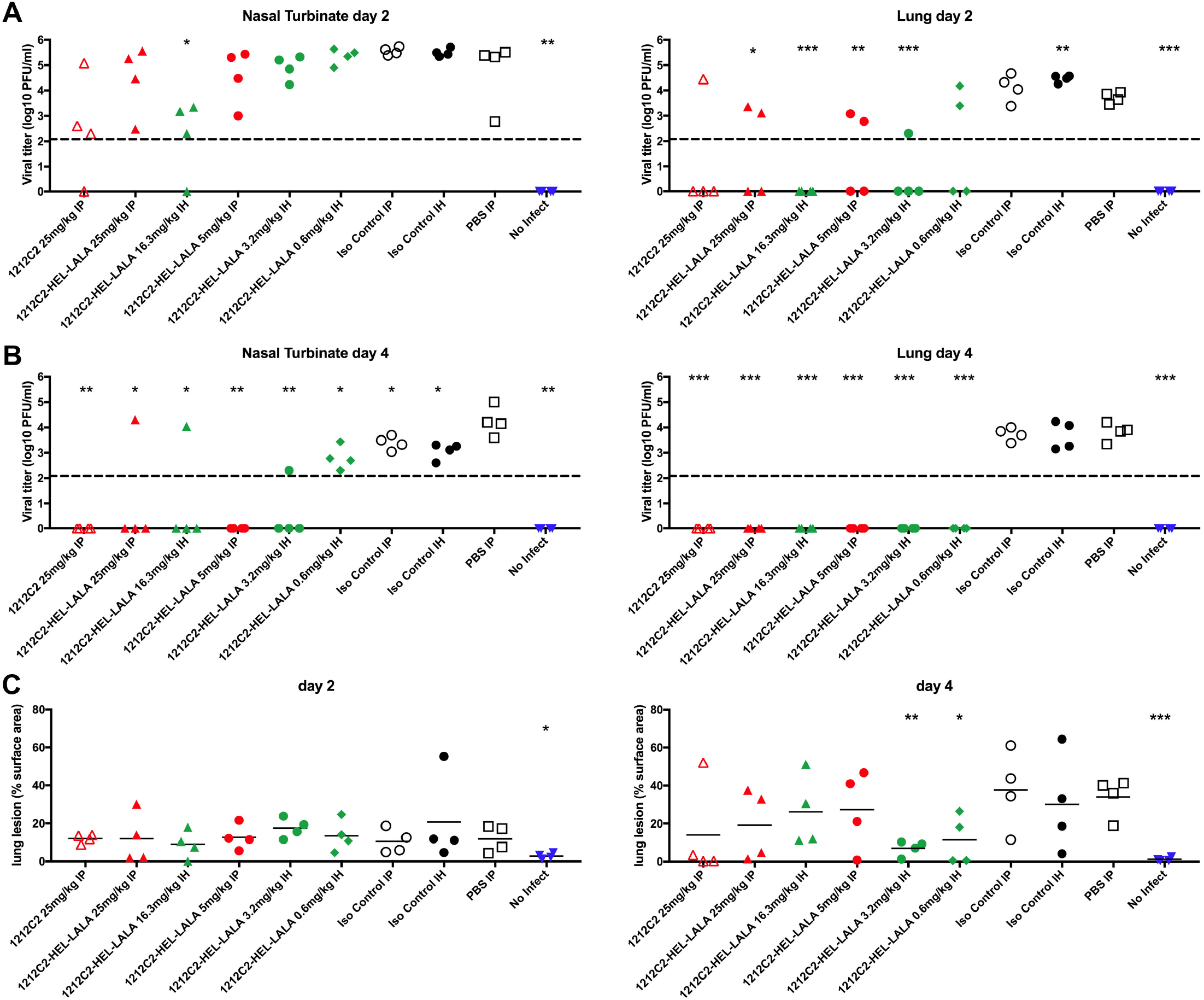
Therapeutic activity of inhaled 1212C2 hmAb in SARS-CoV-2 infected hamsters. Golden Syrian hamsters were infected intranasally with 2×10^5^ PFU SARS-CoV-2 and 12h later treated with indicated hmAb by IP or IH administration. Virus present in nasal turbinates and lungs was determined by plaque assay at 2 dpi (**A**) and 4 dpi (**B**). Dotted line indicates limit of detection. (**C**) Distribution of pathologic lesion were measured using ImageJ and represented as the percent of the total lung surface area. * p<0.05, ** p<0.005, *** p<0.0005 compared to PBI IP treated group.

At 4 dpi virus was detected in the nasal turbinate and lungs of all infected control hamsters (saline and isotype hmAb), with a trend toward modest non-specific reduction in viral titer in nasal turbinate only with isotype control hmAb (**Fig 6B**). Viral titers were only sporadically detected in the nasal turbinates of the 1212C2 treated groups, with the exception of detectable virus in all of the 0.6 mg/kg inhaled 1212C2-HLE-LALA group. At 4 dpi there was no detectable viral titer in the lungs of any 1212C2 treated hamsters, including the 0.6 mg/kg IH 1212C2-HLE-LALA group, representing at least a 2-log reduction compared to control groups. Lung pathology was evident at 4 dpi, particularly in the control groups, however overall, relative to non-infected hamsters, no lung pathology was evident in 9/24 (37.5%) of the 1212C2 treated hamsters, compared with lung pathology evident in 11/12 (91.7%) of the control treated infected hamsters (**Fig 6C**). At 4 dpi lung lesions were significantly decreased in hamsters that were treated with 3.2 mg/kg and 0.6 mg/kg of inhaled 1212C2 hmAb compared to PBS treated hamsters.

With regard to effector function removal brought about by the addition of the LALA mutation, animals treated 1212C2-HLE-LALA appeared to achieve comparable viral reduction as the un-modified 1212C2 mAb. Overall these results confirm the therapeutic activity of 1212C2 hmAb against SARS-CoV-2 infection and suggest increased efficacy of inhaled 1212C2 hmAb, with elimination of viral lung burden with a single inhaled dose of just 0.6 mg/kg. As shown in **Suppl. Table 2**, the lung deposited dose in hamsters is approximately 1.7% of the inhaled dose. This implies that elimination of lung viral burden was achieved at day 4 post-infection with a lung dose of 0.01 mg/kg. Further, if the relevant translation between species to achieve an efficacious dose is a normalization based on a lung-deposited dose per kilogram of lung weight, then, combined with the expectation that the at least 50% of the inhaled dose from a similar commercial nebulizer would be deposited in the lung of a human, the human equivalent efficacious inhaled dose would be 0.03 mg per kg body weight (assumptions: 110g hamster with 1 gram lung weight; 70 kg human with 1000g lung weight).

## Discussion

The SARS-CoV-2 global pandemic continues without clear evidence yet of an effective vaccine to prevent infection or optimal targeted intervention to treat the infection. Our results clearly demonstrate that SARS-CoV-2 infection can result in memory B cells encoding high affinity, high potency NAbs specific for the RBD. The 1212C2 hmAb is able to significantly reduce viral burden in SARS-CoV-2 infected hamsters when used either prophylactically or therapeutically. The *in vitro* affinity compared favorably to and the neutralization activity of 1212C2 was similar to hmAbs that are in late stage clinical trials to treat SARS-CoV-2, suggesting that 1212C2.

The resulting panel of SARS-CoV-2 neutralizing hmAbs indicate that RBD-specific IgG, IgA, and IgM memory B cells develop after infection, however, exhibit only modest somatic hypermutation, which is consistent with an acute primary immune response. Several hmAbs, including 1212C2, 1212F5, 1213H7, and 1212F2 despite only modest somatic hypermutation (4.1-8.2% VH) have remarkably high affinity for SARS-CoV-2 RBD (104-1180 pM *K*D) and potent neutralization (<200 ng/ml NT_50_). The frequent usage of VH3-66 and Vk1-9 among the hmAbs may represent preferential RBD specificity among those germline genes. 1212C2 and 1212F5, which are members of the same clonal lineage, were isolated from IgM memory B cells, and utilize the VH1-2 heavy chain gene. VH1-2 usage is a common feature of other antibodies targeting viral glycoproteins, including HIV envelope (*27*–*29*). VH1–2 usage is more pronounced among marginal zone B cells than naive B cells or switched memory B cells and increased among splenic marginal zone B cell lymphoma (*30*–*33*). This may indicate that the 1212C2 lineage arose from a marginal zone B cell response.

1212C2 has higher affinity (*K*D =104 pM) for SARS-CoV-2 RBD compared to 1212F5 (*K*D =841 pM) which is consistent with its further somatic hypermutation. Additional lineage members were identified, with greater somatic hypermutation, and may be used to guide rational improvement of 1212C2 affinity, neutralization, and ultimate efficacy against SARS-CoV-2 infection.

The resulting hmAbs all neutralized SARS-CoV-2 at NT_50_ of 1 μg/ml or less, indicating their utility for investigating the mechanisms and epitopes mediating SARS-CoV-2 RBD targeted anti-viral activity. Although there was overall consistency that the hmAbs neutralized SARS-CoV-2 either when pre-incubated with the virus (pre-treatment) or after the cells were infected with the virus for 1 h (post-treatment), particularly 1212C2, 1212F5, and 12137 potently neutralized (NT_50_<100 ng/ml) SARS-CoV-2 in both conditions, discordance was noted. 1206D1, which was tested *in vivo*, was approximately 2-fold less effective neutralizing in post-treatment than pre-treatment, which may contribute to its limited *in vivo* activity. In contrast, 1215D1, which had the highest affinity for RBD (*K*D = 73 pM) is approximately 5-fold more effective in neutralizing in post-treatment than pre-treatment. It is expected that the post-treatment neutralization assay may identify antibodies that are better able to mitigate cell-to-cell virus spreading, including antibodies targeting the fusiogenic activity of the S2 domain. Subsequently, 1215D1 should be evaluated for its *in vivo* efficacy, and discern if the post-treatment neutralization assay is a more sensitive indicator of *in vivo* efficacy. Epitope mapping suggests that 1212C2 has a large footprint on the RBD, blocking the binding of several other neutralizing hmAbs encoded by diverse heavy and light chain variable region genes. Efforts are ongoing to solve the 1212C2 – RBD structure to adequately define the epitope.

The high affinity binding of 1212C2 to SARS-CoV-2 RBD, mediating its ability to block RBD attachment to ACE2, and subsequent potent neutralization of SARS-CoV-2 even when added after virus has been added to culture, demonstrate its direct and substantial anti-viral activity. These properties likely contribute to its prophylactic and therapeutic ability to protect hamsters from SARS-CoV-2 challenge, reducing viral burden and the development of lung pathology. 1212C2 was similarly effective in treating SARS-CoV-2 infection in hamsters, reducing viral burden and lung pathology when administered by parenteral route at clinically relevant doses.

As the portal of entry for SARS-CoV-2 virus is the respiratory tract, with the lungs serving as the key target organ for pathogenesis, delivering directly the mAbs to the lungs using inhalation is a logical approach. While aerosols delivery of drugs to the lungs is significantly more inefficient in small animals as compared to humans (~1% of the inhaled dose is deposited in the lungs for rodents as compared to ~50% in humans), our data showed that inhaled delivery of mAbs to the lung BAL in hamsters is still substantially more efficient than that achieved using the IP route (**Supplemental Table 2**), When comparing the inhaled route to the IP route in the hamster challenge study, therapeutic efficacy was achieved at a significantly higher efficiency was observed for the inhaled route. Inhaled administration of 1212C2 hmAb resulted in sterilizing therapeutic protection at all tested doses, with the lowest inhaled dose of 0.6 mg/kg which corresponds to 0.01 mg/kg of lung deposited dose (or a human equivalent inhaled dose of 0.03 mg/kg). In contrast, mAbs against SARS-CoV-2 that have been reported to date required at least 5 mg/kg or above to achieve therapeutic efficacy when administered parenterally (*15, 34–36*). Therefore, inhaled 1212C2 has the potential to profoundly facilitate dose-sparing and treatment coverage compared to conventional parenteral administration. There are several commercial inhaled protein therapeutic products and many more inhaled protein therapeutics that are in various stages of clinical evaluations, most with attractive safety profile. Several mAbs have also been clinically evaluated as inhaled aerosols with demonstrated preliminary safety and tolerability (*6, 16, 37, 38*), suggesting the practicality of formulating 1212C2 for self-administered inhalation. Over 90% of all symptomatic COVID-19 patients are not hospitalized, but they all still need treatment to minimize the potential for transmission and limit complications from viral infection. This is a large unmet COVID-19 population and having a convenient self-administered dose that can be done on an outpatient basis or at home using commercially available nebulizer devices could materially impact treatment coverage, reducing transmissibility, and ultimately the viral burden of the population.

## Materials and Methods

### Sample Collection and B cell isolation

Peripheral blood was collected at the University of Alabama at Birmingham from adult convalescent patients approximately 1 month following PCR confirmed infection with SARS-CoV-2. The patients were symptomatic for at least 7 days and were not hospitalized during SARS-CoV-2 infection. Peripheral blood mononuclear cells (PBMC) were isolated by density gradient centrifugation and cryopreserved. The subjects provided signed written informed consent. All procedures and methods were approved by the Institutional Review Board for Human Use at the University of Alabama at Birmingham, and all experiments were performed in accordance with relevant guidelines and regulations. SARS-CoV-2 recombinant protein was generated, briefly SARS-CoV-2 Spike RBD domain consisted of residues Thr-333 to Thr-531 (uniprot P0DT2). The RBD sequence was cloned in frame with a his8 and avi-tag sequence, respectively (RBDhisavi). RBDhisavi was expressed in insect cells and purified by nickel affinity chromatography. Purified RBDhisavi was biotinylated *in vitro* using BIRA (Avidity.com) for subsequent formation of RBDhisavi-tetramers for B-cell staining experiments. The same RBD sequence was cloned in frame with a murine IgG1 FC domain (RBD-FC). RBD-FC was expressed in insect cells and purified by protein A affinity chromatography. Cryopreserved cells were thawed and stained for flow cytometry similar as previously described (*39*), using anti-CD 19-APC-Cy7 (SJ25C1, BD Biosciences), HIV gp140-AlexaFluor647, SARS-CoV-2 RBD-BV421, CD3-BV510 (OKT3, Biolegend), CD4-BV510 (HI30, Biolegend), IgD-FITC (IA6-2, BD Biosciences), CD27-PE (CLB-27/1, Life Technologies), Annexin V-PerCP-Cy5.5 (Biolegend), and Live/Dead aqua (Molecular Probes).

### Monoclonal antibody production

Single B cells were sorted using a FACSMelody (BD Biosciences) into 96-well PCR plates and immediately frozen at −80 °C until thawed for reverse transcription and nested PCR performed for IgH, Igλ, and Igκ variable gene transcripts as previously described (*27*, *39*). Paired heavy and light chain genes cloned into IgG1 expression vectors and were transfected into HEK293T cells and culture supernatant was concentrated using 100,000 MWCO Amicon Ultra centrifugal filters (Millipore-Sigma, Cork, Ireland), and IgG captured and eluted from Magne Protein A beads (Promega, Madison, WI) as previously described (*27*, *39*). The hmAb 1069D6 or a human myeloma IgG1 (BioXcell, Labanon, NH) were used as an isotype control. 1212C2 was also generated with the incorporation of the LALA mutation to diminish Fc receptor binding (*40*) and a mutation described to increase halflife (*26*). Immunoglobulin sequences were analyzed by IgBlast (www.ncbi.nlm.nih.gov/igblast) and IMGT/V-QUEST (http://www.imgt.org/IMGT_vquest/vquest) to determine which sequences should lead to productive immunoglobulin, to identify the germline V(D)J gene segments with the highest identity, and to scrutinize sequence properties. REGN 10987, REGN 10933, CB6/JS016, and CR3022 were previously described (*15, 17, 41*) and heavy and light chain variable regions synthesized by IDT based on reported sequences (MT470197.1, MT470196.1, DQ168569.1, DQ168570.1 and (*17*)) and cloned into IgG1 expression vector for production in HEK293T cell.

### Binding characterization

ELISA plates (Nunc MaxiSorp; Thermo Fisher Scientific, Grand Island, NY) were coated with recombinant SARS-CoV-2 proteins at 1□μg/ml. Recombinant proteins used include SARS-CoV-2 S1+S2 (Sino Biological, Wayne, PA), SARS-CoV-2 D614G S1 (Sino Biological), SARS-CoV-1 S (BEI Resources), SARS-CoV-2 Nucleocapsid (Sino Biological), and HepG2 whole cell lysate (Abcam, Cambridge, MA). Purified hmAbs were diluted in PBS, and binding was detected with HRP-conjugated anti-human IgG (Jackson ImmunoResearch, West Grove, PA). In selected ELISAs, 8M urea were added to the ELISA plate and the plates incubated for 15 min at room temperature prior to washing with PBS plus 0.05% Tween20 and detection with anti-IgG-HRP to evaluate avidity. Immunofluorescence assay was used to determine hmAb binding to SARS-CoV-2 infected cells. Briefly, confluent monolayers of Vero E6 cells were mock infected or infected with the indicated SARS-CoV-2. At 24 hours post infection (hpi), cells were fixed with 4% paraformaldehyde (PFA) for 30 minutes and permeabilized with 0.5% Triton X-100–PBS for 15 min at room temperature, and blocked with 2.5% Bovine Serum Albumin at 37 °C for 1 h. Cells were then incubated for 1□h at 37°C with 1□μg/ml of indicated hmAb. Then, cells were incubated with fluorescein isothiocyanate (FITC)-conjugated secondary anti-human Ab (Dako) for 1□h at 37 °C. Images were captured using a fluorescence microscope and camera with a 10X objective.

### Surface Plasmon Resonance (SPR)

SPR experiments were performed on a Biacore T200 (Cytiva) at 25°C using a running buffer consisting of 10mM hepes, 150mM NaCl, 0.0075% P20, and 100μg/mL BSA. For affinity measurements, antibodies were captured onto CM-5 sensor chips using the human IgG capture kit (Cytiva). All SPR affinity experiments were double referenced (e.g. sensorgram data was subtracted from a control surface and from a buffer blank injection). The control surface for all experiments consisted of the capture antibody. Approximately 100RU of each hmAb was captured onto the chip surface. RBD was injected over the NmAbs at four concentrations (80nM, 20nM, 5nM, and 1.25nM) for 90 seconds, followed by 300 second dissociation phase. After each injection, the surface was regenerated with a 30 second pulse of 3M MgCl2, followed by capture of fresh hmAb. The buffer flowrate for affinity analysis was 50μL/min. Sensorgrams were globally fit to a 1:1 model, without a bulk index correction, using Biacore T-200 evaluation software version 1.0.

Epitope mapping was performed by capturing RBD-FC to CM-5 sensor chips using an anti-murine FC capture kit (Cytiva). The running buffer was the same as used for RBD / hmAb affinity measurements. hmAbs were sequentially injected, up to 6 NmAbs in one series, over the RBD-FC surface. The first hmAb was injected at 100 nM concentration, and subsequent hmAbs were injected at 50nM concentrations. Each hmAb was injected for 90 seconds over the RBD-FC, followed by a 60 second dissociation phase. The flowrate for the epitope mapping studies was 30 μL/min. After the final injection, the surface was regenerated by a 3-minute injection of 10 mM glycine pH 1.7. hmAb epitopes were defined by measuring the reduction in hmAb binding (RU) to the RBD-FC surface in the presence of different hmAbs, relative to hmAb-RBD-FC alone.

### ACE2 – RBD inhibition assay

All dilutions and cell suspensions were made in MACs buffer [autoMACs rinsing solution (Miltenyi Biotec, Auburn, CA) with 0.5% BSA]. For 20 μg of hmAb, 0.2 μg of biotinylated SARS-CoV-2 RBD was mixed in a total volume of 1 ml and incubated for 30 minutes on ice. HEK293T cells co-expressing human ACE2 and green fluorescent protein (BEI Resources) were grown in DMEM containing 4 mM L-glutamine, 4500 mg per L-glucose, 1 mM sodium pyruvate and 1500 mg per mL sodium bicarbonate, supplemented with 10% fetal bovine serum. On the day of experiment, cells were dissociated with 0.5 mM EDTA and a cell scraper. Cells were washed with MACs buffer and pelleted at 350 x g for 5 minutes. Cells were counted, distributed at 5 x 10^5^ per tube, and pelleted. Cells pellets were suspended in the mAb – RBD mixture and allowed to interact for 60 m on ice. Unbound RBD was washed away by adding 1 ml MACs buffer and centrifuging for 5 minutes at 350 x g. Supernatant was carefully removed, and cells suspended in 100 μl MACs buffer. 7AAD and Streptavidin-Pacific Blue (Invitrogen) was added and the mixture incubated on ice for 30 minutes. To wash cells, 1 ml of MACs buffer was added and tubes centrifuged at 350 x g for 5 minutes. Supernatant was removed and the wash step was repeated with 2 ml of ice cold PBS. After removing the PBS wash, the cell pellet was suspended in 0.5 ml PBS and the cells passed through a cells strainer on a 5 ml tube (Falcon). Cells were analyzed on a FACSymphony (BD Biosciences) and gated on 7AAD negative GFP positive cells.

### SARS-CoV-2 neutralizing activity

hmAbs were tested for neutralization of SARS-CoV2 (isolate USA-WA1/2020, BEI Resources) as previously described (*42*). For pretreatment, 100-200 plaque forming units (PFU)/well of SARS-CoV-2 containing 2-folds dilutions (starting concentration 25 μg/ml) for hmAbs (or 1:100 dilution for human serum control) were mixed in 100 μl of media and incubated at 37 °C for 1h. Vero E6 cells (96-well plate format, 4 x 10^4^ cells/well, quadruplicate) were infected with virus-hmAb or virus-serum (control well) mixture for virus adsorption for 1h, followed by changing media with post-infection media containing 2% FBS and 1% Avicel. For post-treatment, Vero E6 cells (96-well plate format, 4 x 10^4^ cells/well, quadruplicate) were infected with 100-200 PFU/well of SARS-CoV-2. After 1 h of viral adsorption, the infection media was changed with the 100 μl of media containing 1% Avicel and 2-fold dilutions, starting at 25 μg/ml of hmAb (or 1:100 dilution for human serum control). At 24 hpi, infected cells were fixed with 10% neutral formalin for 24 h and were immune-stained using anti-NP monoclonal 1C7C7 antibody (*21*). Virus neutralization was evaluated and quantified using ELISPOT, and the percentage of infectivity calculated using sigmoidal dose response curves. The formula to calculate percent viral infection for each concentration is given as [(Average # of plaques from each treated wells – average # of plaques from “no virus” wells)/(average # of plaques from “virus only” wells - average # of plaques from “no virus” wells)] x 100. A non-linear regression curve fit analysis over the dilution curve can be performed using GraphPad Prism to calculate NT_50_. Mock-infected cells and viruses in the absence of hmAb were used as internal controls. hmAbs were also tested using a SARS-CoV-2 Spike protein pseudotyped virus (PsV) containing the gene for firefly luciferase. Virus neutralization can be measured by the reduction of luciferase expression. VeroE6/TMPRSS2 cells were seeded at 2 x 10^4^ cells/well in opaque plates (Greiner 655083). The next day, PsV corresponding to 1-10 x 10^6^ luciferase units was mixed in Opti-MEM with dilutions of hmAbs and incubated at RT for 1 h. Media was removed from the cells and 100 μl/well of the hmAb/PsV mix was added in triplicates. After 1 h incubation at 37C and 5% CO_2_, another 100 μl of media containing 2% FBS, was added, and cells were incubated for 24 more hrs. After this time, luciferase activity was measured using Passive Lysis Buffer (Promega E1941) and Luciferase substrate (Promega E151A) following the manufacturer’s instructions. Neutralization was calculated as the percent reduction of luciferase readings as compared to no-antibody-controls.

### Prophylactic and therapeutic protective activities of RBD specific hmAbs in hamsters

We have previously demonstrated that golden Syrian hamsters (*Mesocricetus auratus*) are susceptible to SARS-CoV2, showing loss of body weight and viral replication in the lungs and nasal turbinate (*22*). In a prophylactic experiment, 6-week old, female hamsters were IP injection of 10 mg/kg of hmAb 1212C2 (n=6), 1206D1 (n=3), human IgG1 Isotype control mAb (BioXcell, Labanon, NH; n=6), or PBS (n=5). Six hours later, hamsters were given 2×10^5^ PFU of SARS-CoV2, intranasally (IN). Three animals were also mock infected with PBS at this time to serve as a negative control. For each day, lungs and nasal turbinate were collected removed and photographed on white paper towels to look for gross pathology using a macroscopic pathology scoring analysis measure the distributions of pathological lesions, including consolidation, congestion, and pneumonic lesions using ImageJ software (NIH). Resulting pathology were represented as the percent of the total lung surface area. Left side of the lungs were used for histopathology and the right side was used for viral titers.

The hmAb 1212C2 was also assessed in an *in vivo* therapeutic experiment. At the start of the experiment, 12, 6-week old, female hamsters were administered 2×10^5^ PFU of SARS-CoV2, intranasally. Four animals were mock infected with PBS at the same time. Six hours later, animals were given either 25 mg/kg hmAb 1212C2 (n=4), 25 mg/kg isotype control (n=4), or PBS (n=4) IP. Animals were euthanized at 4 dpi and lungs and nasal turbinates collected for histopathology and viral titers as described for the prophylactic experiment.

### Delivery of inhaled 1212C2

The therapeutic testing of 1212C2 was tested using 5-week old, female hamsters. Hamsters were challenged with 2×10^5^ PFU of SARS-CoV2 IN and 12 hours later treated with hmAb IP or IH. The estimated inhaled dose of hmAb is measured by sampling the aerosolized atmosphere the hamsters are exposed at a known fixed flowrate during the entire course of their exposure. The gas drawn out of the exposure chamber by vacuum passes through a collection filter to capture the aerosolized droplets. The amount of mAb deposited in the filter is determined using A280 following extraction in formulation buffer, which determines the mAb aerosol concentration in the exposure chamber. The approximate inhaled dose over a 30 minutes whole body exposure is then derived from the aerosol concentration and minute volume of the hamsters. The target high dose level of nonspecific IgG or 1212C2-HLE-LALA was 25 mg/kg. The actual dose measured was 11.30 mg/kg for the non-specific IgG group and 16.3 mg/kg for the 1212C2-HLE-LALA group (**Table 2**).

**Table 2.** Estimated inhaled 1212C2 dose

|  | Nebulizer<br>Consumption<br>Rate | Aerosol<br>Conc. | Aerosol<br>mAb<br>Conc. | Dose | Actual<br>Dose<br>(ave.) |
| --- | --- | --- | --- | --- | --- |
| Description | mL/min | mL/L | mg/L | mg/kg | mg/kg |
| Nonspecific IgG Inhalation 25 mg/kg (cage 1) | 0.30 | 0.033 | 0.51 | 9.13 | 11.30 |
| Nonspecific IgG Inhalation 25 mg/kg (cage 2) | 0.25 | 0.049 | 0.75 | 13.48 |  |
| No Virus, Saline Inhalation (cage 1) | 0.19 | Not<br>measured | 0.00 | 0.00 | 0.00 |
| No Virus, Saline Inhalation (cage 2) | 0.17 | Not<br>measured | 0.00 | 0.00 |  |
| 1212C2 HLE-LALA Inhalation 25 mg/kg (cage 1) | 0.38 | 0.052 | 0.81 | 14.52 | 16.33 |
| 1212C2 HLE-LALA Inhalation 25 mg/kg (cage 2) | 0.38 | 0.065 | 1.01 | 18.15 |  |

The mid and low dose groups for 1212C2-HLE-LALA were 5-fold and 25-fold dilutions of the mAb solution used in the high dose, respectively, which correspond to 3.2 mg/kg and 0.6 mg/kg.

### Deep-sequencing immunoglobulin repertoire analysis

10 million PBMCs were thawed and used for RNA isolation (Qiagen, RNeasy Mini Kit). Removal of any residual genomic DNA was performed using the Turbo DNA-Free kit (Invitrogen) and cDNA was generated using the qScript cDNA synthesis kit (QuantaBio). Two rounds of PCRs were performed to generate the libraries with either global (VH1 – VH6) or individual VH targeted forward primers as previously described (*27*, *39*) with the modifications adapted for Illumina Nextera approach. The final PCR products were gel extracted (QIAQuick, Qiagen, Hilden, Germany) and further purified using the ProNex size-selective purification system (Promega) to select products between 500-700 bp range. Libraries were submitted to the Heflin Center for Genomic Sciences at the University of Alabama at Birmingham where DNA quantification was made by qPCR and sequenced on an Illumina MiSeq system (Illumina, Inc., CA, USA) using 2 × 300 bp paired-end kits (Illumina MiSeq Reagent Kit v3). Sequence analysis and assembly of lineage trees were performed using an in-house custom analysis pipeline as previously described (*39*, *43*). All sequences were aligned using IMGT.org/HighVquest (*44*). Lineage trees were generated by identifying the lineage (the cluster of sequences with identical VH, JH, and HCDR3 lengths and ≥85% HCDR3 similarity) containing the corresponding hmAb sequence.

### Statistical analysis

Significance was determined using GraphPad Prism, v8.0. *t* tests were applied for evaluation of the results between treatments. For statistical analysis viral titers were log transformed and undetectable virus was set to the limit of detection (200 PFU/ml).

## Acknowledgements

We are grateful for the clinical research staff that enabled this project and for the technical assistance provided by Christopher Bates, Reuben Burch, Eric Carlin, and Sarah Sterrett, and the assistance of the University of Alabama at Birmingham Center for AIDS Research and its Flow Cytometry Core Facility under the direction of Olaf Kutsch, and the UAB Heflin Genomics Core, UAB Multidisciplinary Molecular Interaction Core Facility. We thank Alexander Rosenberg and Christopher Fucile for providing immunoglobulin sequence analysis software. We are most grateful for the participation of the study volunteers.

