## Supplemental Table 1 for "Therapeutic activity of an inhaled potent SARS-CoV-2 neutralizing human monoclonal antibody in hamsters"

| hmAb | $NT_{50}$ (ng/ml) |
|---|---|
| 1212C2 | 1.9 |
| 1212F5 | 3.5 |
| 1213H7 | 2.9 |
| 1215D1 | 18.8 |
| 1212D5 | 38.7 |
| 1212F2 | 58.3 |
| SAD35 | 457 |
| REGN10987 | 4.5 |
| CB6/JS016 | 114 |

**Supplemental Table 1. SARS-CoV-2 Pseudovirus neutralization assay**
