## Supplemental Table 2 for "Therapeutic activity of an inhaled potent SARS-CoV-2 neutralizing human monoclonal antibody in hamsters"

**Supplemental Table 2.** Hamsters (N=2 per group) were treated using a hmAb at approximately 25mg/kg by intraperitoneal or inhalation route as indicated. Serum, lung and bronchial lavage (BAL) samples were collected at 30 mins or 42 hours after each treatment from each animal. The titer of hmAb recovered from each sample was quantitated using anti-human IgG ELISA assay.

| **Inhaled Group (IH)** | **30 minutes post-admin** | **42 hours post-admin** |
|---|---|---|
| Nebulizer output rate (mL/min) | 0.40 | 0.43 |
| Aerosol liquid concentration (mL/L) | 0.08 | 0.10 |
| **Inhaled dose (mg/kg)** | **26.5** | **32.0** |
| Serum conc (ng/mL) | 102 | 872 |
| Total in serum (ng) [78 mL blood/kg, hematocrit 43%] | 453 | 3877 |
| **% of inhaled dose in serum** | **0.017** | **0.121** |
| Total in BAL (ng) | 44980 | 6032 |
| **% of inhaled dose in BAL** | **1.699** | **0.189** |

| **Intraperitoneal Group (IP)** | **30 minutes post-admin** | **42 hours post-admin** |
|---|---|---|
| **Estimated dose (mg/kg)** | **26.9** | **26.9** |
| Serum conc (ng/mL) | 2986 | 17722 |
| Total in serum (ng) [78 mL blood/kg, hematocrit 43%] | 13275 | 78791 |
| **% of delivered dose in serum** | **1.38** | **7.03** |
| Total in BAL (ng) | 0.000 | 287 |
| **% of delivered dose in BAL** | **0.000** | **0.026** |
